# *Psychromonas aestuarii* sp. nov., a novel bacterial species isolated from estuarine surface sediment

**DOI:** 10.1101/2025.01.26.634906

**Authors:** Iuliana Nita, Yaren Kart, Mikael Lenz Strube, Mikkel Bentzon-Tilia

**Affiliations:** Department of Biotechnology and Biomedicine, Technical University of Denmark, Søltofts Plads 221, Lyngby, Denmark

**Keywords:** Estuary, iChip, novel species, *Psychromonas*, *Psychromonas aestuarii*, sediment

## Abstract

Using an isolation chip (iChip), two bacterial strains, MME1^T^ and MME2, exhibiting ovoid to curved rod cell morphologies and white-cream colony pigmentation were isolated from the surface of estuarine sediment at Nivå Bugt Strandenge Bird Sanctuary, Denmark (55°55’42.6’N 12°31’23.6’E). Analysis of the 16S rRNA gene sequence of these isolates suggested them to be members of the genus *Psychromonas*, as both had above 97.5 % 16S rRNA gene similarity with *Psychromonas aquimarina strain* JAMM 0404^T^. Further genomic comparisons suggested MME strains to be representatives of a novel species, having an average nucleotide identity (ANI) < 85 % relative to all other genome sequenced species of the *Psychromonas* genus. The G+C content was 39.5 mol %. The species was facultative anaerobic, growing optimally at 15 – 20 °C and pH 7.0. The optimal salinity was lower than for other described species of the genus with 1 – 2 % NaCl, reflecting the estuarine source of isolation. GC-MS analyses identified C_16:0_, C_16:1_^ω7c/^ C_16:1_^ω7t/^ and C*_14:0_^3OH^* as the predominant cellular fatty acids. Based on molecular, phenotypic, and chemotaxonomic analyses, we propose that MME1 and MME2 represent a novel species of the genus *Psychromonas* with the name *Psychromonas aestuarii* sp. nov. with the MME1^T^ (= DSM 118464^T^ =LMG 33722^T^) as the type strain.

## INTRODUCTION

The genus *Psychromonas* was first described in 1998 with the identification of *Psychromonas antarctica* (1), the type species of the genus. Currently there are 15 described *Psychromonas* spp., with the most recently characterized species being *Psychromonas aquatilis* M1A1^T^, isolated from Antarctica in 2017 (2). The genus *Psychromonas* is the only genus of the family *Psychromonadaceae*, which relate most closely to *Pseudoalteromonadaceae*, *Colwelliaceae*, and *Idiomarinaceae* (3). Members of the genus *Psychromonas* are Gram-negative with rod or ovoid cells, which are usually, but not exclusively, motile (3). They typically have a G+C content ranging between 38 and 43.8 mol % and generally exhibit psychrophilic and halophilic traits, with membranes rich in saturated C_16:0_ and unsaturated C*_16:1_^ω7^* fatty acids. A few representatives also synthesize polyunsaturated fatty acids (4). The cell size, temperature range, piezophily, presence of gas vacuole and carbon source utilization vary among *Psychromonas* spp. (1,2,5–14). These bacteria have commonly been isolated from cold, saline marine environments (3), including marine sediments, suggesting a role as heterotrophic degraders of organic matter in cold and occasionally anoxic sediments as well as on sea ice and in seawater in arctic regions (1,3,8,9).

In this study, we report the isolation and characterization of two bacterial strains, MME1^T^ and MME2, isolated from estuarine sediments at Nivå Bugt Strandenge Bird Sanctuary, Denmark, designated *Psychromonas aestuarii* MME1^T^ and MME2, representing a novel species of the genus *Psychromonas*.

## ISOALTION AND ECOLOGY

Surface sediment samples (depth 1 – 2 cm) were collected in 50 mL Falcon tubes on March 15, 2021, at Nivå Bugt Strandenge Bird Sanctuary (55°55’42.6’N 12°31’23.6’E) (15), Denmark. Samples were transported to the Molecular Microbial Ecology laboratory at the Technical University of Denmark for microbial cell extraction and cultivation alongside 10 kg of sediment and seawater collected in acid rinsed, closed buckets.

Microbial cells were extracted by rigorous mixing of 1 g of sediment in 9 mL of 2 % (w/v) NaCl and sonication at 8 Amp for 3 × 20 s intervals. After 15 min of settling, the supernatant was transferred to a fresh tube and stored at 4 °C overnight. Five mL of 10^-2^ and 10^-3^ dilutions of the cell suspension were filtered through 5 μm polycarbonate and 1 μm cellulose membranes. Bacterial abundances of the cell suspensions were determined by counting cells on the filters using an Olympus BX51 fluorescence microscope after 15 min staining with 20 µL of 10 % SYBR Gold in the dark. Stained filters were mounted on Superfrost slides with PBS containing 43.5 % (v/v) glycerol and 0.0001 % (w/v) p-phenylene-diamine.

Three bacterial cell concentrations (0.5, 1.0, and 10 cells/chamber) in molten ½ MA (1:1, filtered seawater: marine broth, 2216 BD Difco, 2% agar) were loaded into an in-house constructed isolation chip (iChip) for cultivation (16). The iChip was assembled by sealing a membrane (polycarbonate 0.01 μm pore size, Frisenette 1223450) to a sterile pipette tip rack with silicone glue. After drying under UV-light, 48 chambers were inoculated and four were filled with ½ MA medium (negative controls) and the iChip was sealed with a second membrane filter. The iChip was placed on top of a 4 cm layer of sediment from the sampling site in a 40 L aquarium and covered with a 1 - 2 cm sediment layer and a 5 cm layer of seawater. The iChip was incubated at 16 °C for 10 days. The content of the individual chambers was homogenized by passage through a sterile 1 mL syringe into 50 % (v/v) glycerol after which they were stored at −80 °C. Stocks were revived at 16 °C in ½ MB (1:1, filtered seawater: marine broth, BD Difco 2216) and streaked onto ½ MA (1:1, filtered seawater: MB, 2% agar). Individual colonies were picked for further analyses. MME1^T^ and MME2 originated from two different chambers.

## MOLECULAR IDENTIFICATION

Initial taxonomic identification of the MME strains was done by 16S rRNA gene amplification using universal primers 27F (5’-AGAGTTTGATCMTGGCTCAG) and 1492R (5’-TACGGYTACCTTGTTACGACTT) (17) followed by Sanger sequencing at Macrogen (Amsterdam, Netherlands). These sequences were then identified using NCBI blast, which suggested *Psychromonas aquimarina* JAMM 0404^T^ as the closest related species (97.5 %) of both isolates. A phylogenetic tree was generated using MEGA (Molecular Evolutionary Genetics Analysis) v. 11.0.13 (18). 16S rRNA gene sequences of MME strains 1 and 2 were first aligned to the 16S rRNA genes of the 15 previously described *Psychromonas* type strains along with *Idiomarina abyssalis* KMM 227^T^ (19) as the outgroup using CLUSTAL_W. The evolutionary relationships were inferred by the neighbour-joining method (20) building a bootstrap consensus tree (1000 replicates) (21) with distances calculated using the Maximum Composite Likelihood method (22), revealing the placement of MME1^T^ and MME2 as a monophyletic group within the genus *Psychromonas*, with *Psychromonas aquimarina* JAMM 0404T, *Psychromonas ossibalaenae* JAMM 0738^T^ and *Psychromonas macrocephali* JAMM 0738^T^ as closest related species (Fig 1A).

**Fig 1.**
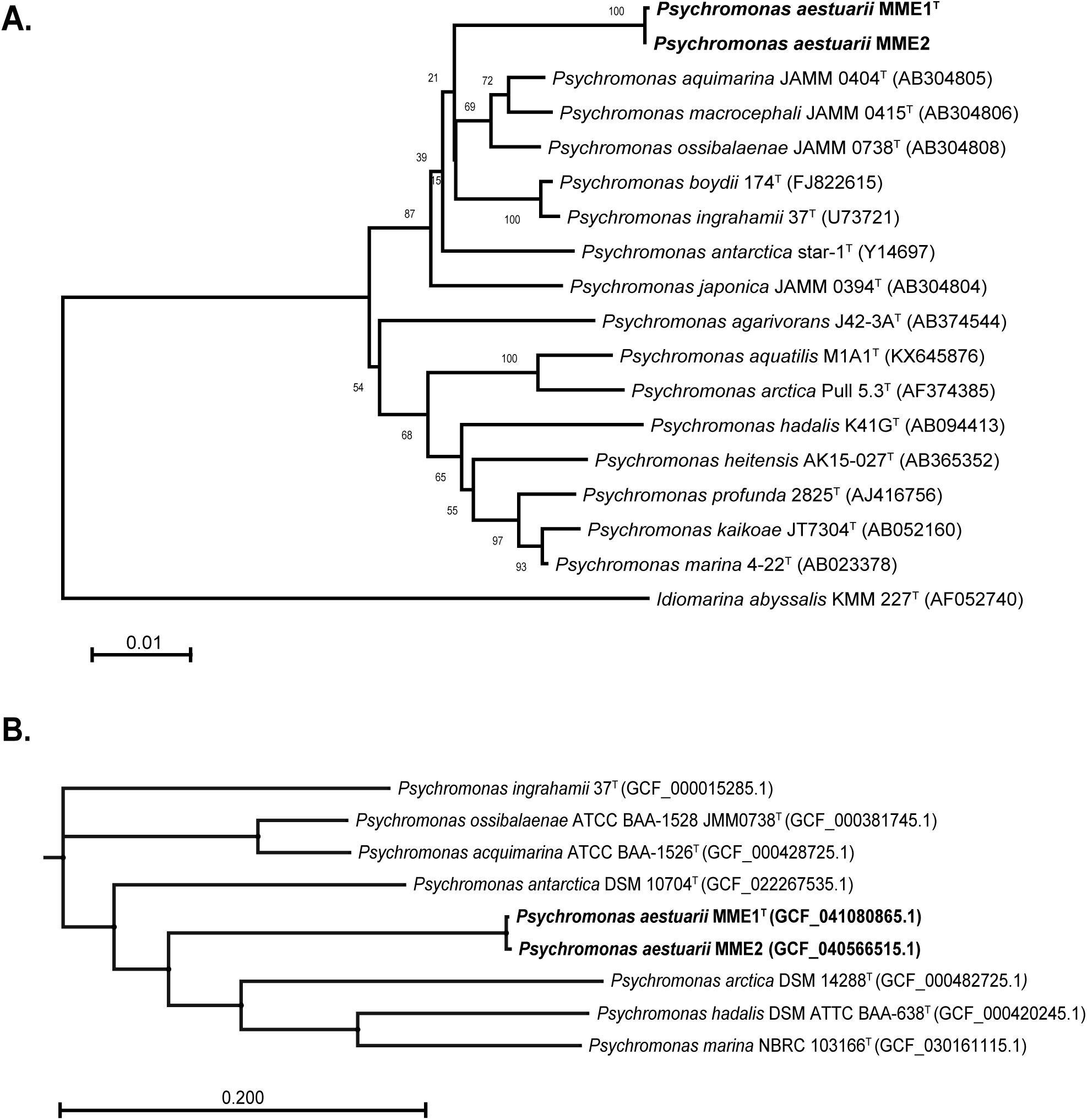
Phylogenetic relationships between *Psychromonas* sp. MME1^T^ and MME2, and type strains of other *Psychromonas* spp. based on 16S rRNA gene sequences **(A)**, and phylogenomic relationships between *Psychromonas* sp. MME1 and MME2, and type strains of *Psychromonas* spp. for which genome sequences were available **(B)**.

To further assess the genetic characteristics and taxonomic classification of MME1^T^and MME2, genomic DNA was extracted using the NucleoSpin Tissue kit (Macherey-Nagel), following the instructions provided by the manufacturer including lysis of cells in 2 % PYMSB (5 g/L Peptone, Bacto; 5g/L yeast extract, Bacto; 3.2 g/L MgSO_4_7H_2_O, Merck) at 56 °C for 18 h. The purified DNA was sequenced on the Oxford Nanopore GridION™ Mk1 platform using the Rapid barcoding kit 24 V14. The resulting sequences were next trimmed for adaptors and filtered for quality before assembly with flye (23) using the dragonflye workflow (https://github.com/rpetit3/dragonflye). The assemblies where then annotated with Bakta (24) and submitted to NCBI as *Psychromonas* sp. MME1 (CP190906) and *Psychromonas* sp. MME2 (JBEWXI000000000). These assemblies where then used for whole-genome comparisons with the 7 existing genomes of *Psychromonas*: First, pairwise average nucleotide identity (ANI) was calculated using MUMMER. Secondly, genes suitable for comparison between all genomes (n=867) were selected and aligned by PANTA, from which a tree was built using FasTree (25).

Genomic comparisons with available complete genomes of the *Psychromonas* genus (n=7) showed that the MME strains are highly similar (ANI > 99.5 %) but have less than 85 % identity to other species (Fig S1). Phylogenomic analysis using 867 core genes further confirmed the close relationship among the MME genomes, while indicating a distant relationship with other *Psychromonas* species (Fig 1B). The G+C content of 39.5 mol% aligned with previously reported data for *Psychromonas* spp.

## MORPHOLOGY, PHYSIOLOGY, AND CHEMOTAXONOMY

Colony morphology was assessed by cultivating MME strains on Marine agar (MA, BD Difco2216) for 2 – 3 days at 20 °C. Colonies were small, < 1 mm, circular with creamy-white pigmentation. They would eventually, over 10 – 14 days grow into lager colonies of similar colour. In marine broth (MB, BD Difco 2216) cultures, growth was slower and reached a maximum optical density of around OD_600_ 0.4. For phenotypic tests, colonies were harvested, suspended in selected media, centrifuged at 1000 rpm for 1 min to remove debris, and ultimately the cleared culture was adjusted to OD_600_ 0.2 for inoculations. Temperature-dependent growth of MME strains was assessed on MA at 2, 5, 10, 15, 18, 20, 22, 25, 27, or 30 °C at day 12. Growth was observed from 2-22 °C with optimal growth at 15 – 20 °C.

Optimal salinity for growth was assessed in PSMSB and PSMSA (PSMSB, 1.5 % agar) supplemented with 0, 1, 2, 3, 4, 5, 6, 7, 8, 9, 10, 11, 12, 13, 14, or 15 % NaCl at 20 °C after 14 days. Growth occurred between 1 – 2 % NaCl on solid substrates and 0 – 3 % in liquid substrate, with an optimal concentration of 1 – 2 % NaCl. Growth was minimal or absent at higher salinities. Optimal pH for growth on PSMSA and PSMSB media, supplemented with 2% NaCl, was assessed for pH 5.0, 5.5, 6.0, 6.5, 7.0, 7.5, 8.0, 8.5, and 9.0 at 20 °C. Growth was observed from pH 6.0 - 7.5 on solid and 6.0 - 9.0 in liquid, with optimal growth at pH 7.0. Motility was assessed at 20 °C for 17 days in semi-solid MA with agar concentrations of 0.3 %, 0.5 %, and 0.7 %. Motility was confirmed in 0.3 % agar between day 7 and 17.

Gram-reaction was determined using 3 % KOH (Ryu method; 26,27). Cells were 1.05 × 1.9 µm and had an ovoid-curved rod morphology. Cytochrome C oxidase was positive using BD BBL Dry Slide Oxidase slides and catalase activity was positive, confirmed by gas formation upon H_2_O_2_ addition to cell pellets. We observed growth of MME1^T^ and MME2 at 15 °C on MA plates after 11 days under anaerobic conditions (Oxoid chamber with Oxoid AnaeroGen Sachets).

API20NE and API50CH tests (bioMérieux) were performed for Gram-negative biochemistry and carbohydrate oxidation at 20 °C for 5 days. MME isolates were resuspended in modified ½ artificial sea water (1.5 % NaCl, 0.035 % KCl, 0.54 % MgCl_2_ · 6H_2_O, 0.27 % MgSO_4_ · 7H_2_O, 0.05 % CaCl_2_ · 2H_2_O; 5) and showed to utilize D-xylose, D-galactose, D-glucose, D-fructose, N-acetylglucosamine, D-cellobiose, and D-maltose with minor variations for MME1^T^ (Table 1). In addition, the strains hydrolyse potassium 5-ketogluconate, esculin and tested positive for β-galactosidase, catalase, cytochrome oxidase, and nitrate reduction.

**Table 1.**
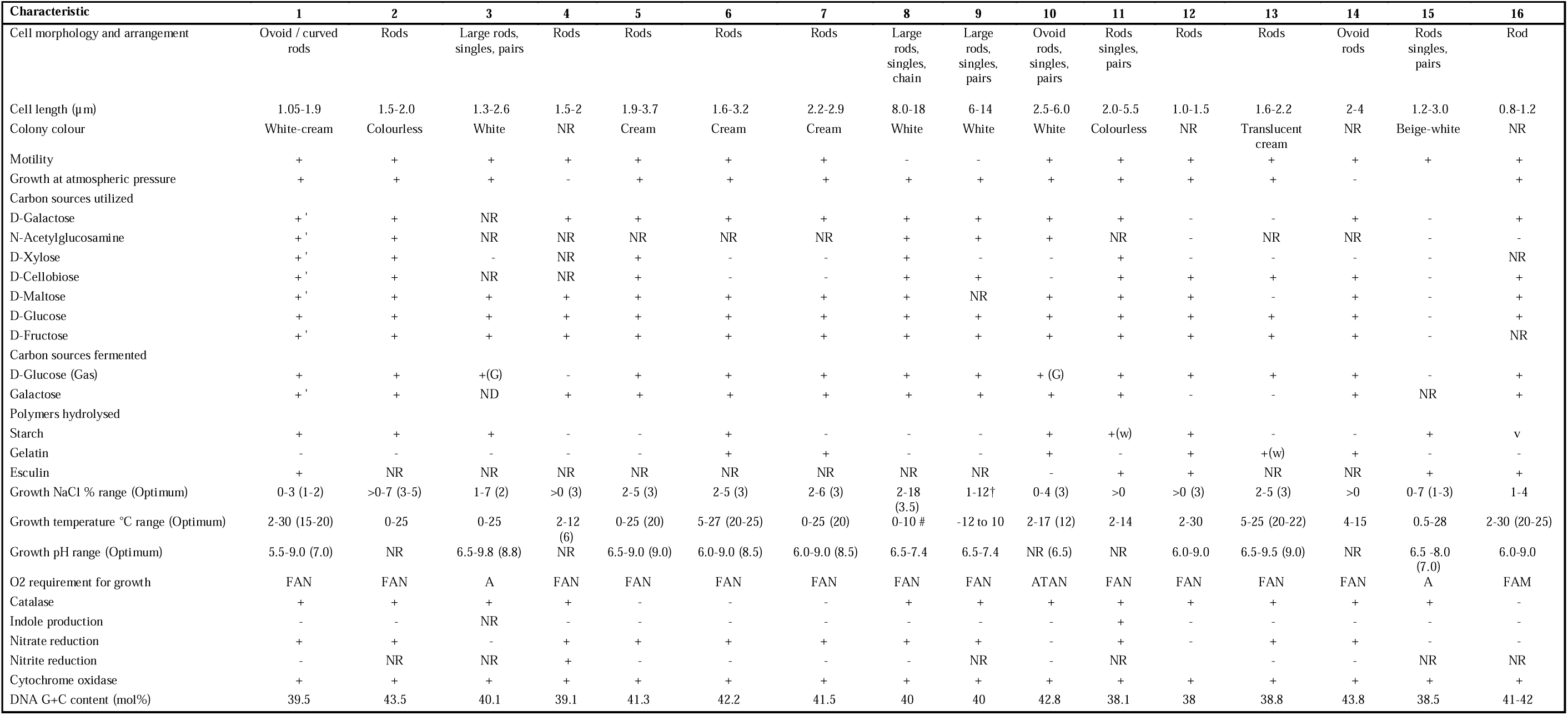
Comparison of characteristics of *Psychromonas aestuarii* MME strains and other members of the genus *Psychromonas*. Strains: 1, *P. aestuarii* MME1^T^ and MME2; 2, *P. marina* 4-22^T^; 3, *P. arctica* Pull 5.3^T^; 4, *P. hadalis* K41G^T^; 5, *P. ossibalaenae* JAMM 0738^T^; 6, *P. aquimarina* JAMM 0404^T^; 7, *P. macrocephali* JAMM 0415^T^; 8, *P. boydii* 174^T^; 9, *P. ingrahamii* 37^T^; 10, *P. antartica* DSM 10704^T^; 11, *P. profunda* 2825^T^; 12, *P. heitensis* AK15-027^T^; 13, *P. japonica* JAMM 0394^T^; 14, *P. kaikoae* JT7304^T^; 15, *P. aquatilis* M1A1^T^; 16, *P. agarivorans* J42-3A^T^; MME data are from this study; other *Psychromonas* species data are from (1,2,5–14). All strains are Gram-negative. Characteristics are scored as: +, positive; -, negative; +‘, positive for one MME strain, but inconclusive for the other; w, weak; NR, not reported; # Growth ≤ 0 °C was not determined; FAN, facultative anaerobe; ATAN, aerotolerant anaerobe; A-aerobe.

Moreover, MME strains can ferment glucose, resulting in acid production. Negative results were observed for nitrite reduction, indole production, arginine dehydrolase, urease, and gelatin hydrolysis. Results revealed physiological differences from type strains of *Psychromonas*, and thereby summarized in Table 1. Complete physiological characteristics of the MME strains are summarized in the species description.

Cellular fatty acid analysis was conducted by Deutsche Sammlung von Mikroorganismen und Zellkulturen (DSMZ) using ∼250 mg of cell pellet obtained from cultures grown in MB for 3 weeks at 15 °C. The pellet was re-suspended in 500 µL isopropanol. Fatty acids were analysed as methyl esters (FAMEs) following previously reported protocols (28), separated by gas chromatography (GC), and identified by GC-MS on an Agilent 7000D system (29).

The cellular fatty acid profile of MME strains aligns with those of *Psychromonas* species (Table 2), featuring predominant saturated fatty acids: C_16:0_ (24.1% MME1^T^, 19.5% MME2), C*_14:0_*^3OH^ (17.4% MME1^T^, 14.5% MME2), *cis*-unsaturated fatty acids C*_16:1_^ω7c^* (13.1% MME1^T^, 19.3% MME2) and *trans*-unsaturated C_16:1_^ω7t^ (24.1% in MME1^T^, 14.1% in MME2). The ability of microbes to convert *cis*-to *trans*-unsaturated fatty acids plays an important role in modulating membrane fluidity in response to external stimuli (30) as reported for the marine *Vibrio* ABE-1 strain. This confers ecological advantages, facilitating rapid adaptation to variations in temperature and salinity (31). Moreover, the MME strains uniquely produce C_13:0_, C_13:0_^3OH^, C*_17:0_*, and C*_18:0_* fatty acids, distinguishing them from other known *Psychromonas* species (Table 2).

**Table 2.**
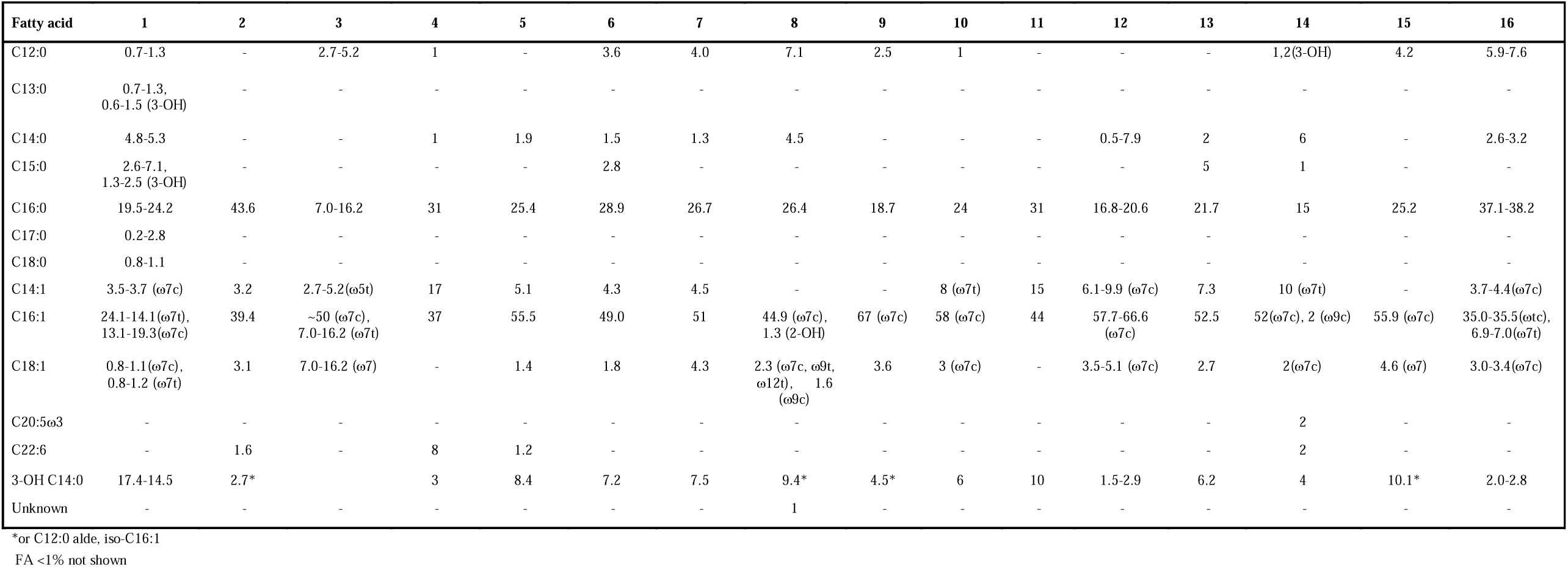
Fatty acid contents of MME strains and other members of the genus *Psychromonas*. Strains: 1, *P. aestuarii* MME1^T^ and MME2; 2, *P. marina* 4-22^T^; 3, *P. arctica* Pull 5.3^T^; 4, *P. hadalis* K41G^T^; 5, *P. ossibalaenae* JAMM 0738^T^; 6, *P. aquimarina* JAMM 0404^T^; 7, *P. macrocephali* JAMM 0415^T^; 8, *P. boydii* 174^T^; 9, *P. ingrahamii* 37^T^; 10, *P. antartica* DSM 10704^T^; 11, *P. profunda* 2825^T^; 12, *P. heitensis* AK15-027^T^; 13, *P. japonica* JAMM 0394^T^; 14, *P. kaikoae* JT7304^T^; 15, *P. aquatilis* M1A1^T^; 16, *P. agarivorans* J42-3A^T^; MME data are from this study; other *Psychromonas* species data are from (1,2,5–14). Values indicate total fatty acid percentages, with isomers in parentheses, if identified. Results < 1 % are not shown, except for MME strains.

Based on genotypic, chemotaxonomic and physiological data, MME strains represent a novel species of the genus *Psychromonas*, for which the name *Psychromonas aestuarii* sp. nov. is proposed.

## DESCRIPTION OF PSYCHROMONAS AESTUARII SP. NOV

*Psychromonas aestuarii* (a.es.tu.a’ri.i. L. n. aestuarium, of the estuary) Cells are Gram-reaction-negative, ovoid-curved rods with a cell size of approximately 1 × 2 µm in size. Colony colour is white-cream and circular after 3 days of incubation on MA at 20°C. The species is motile, facultative anaerobic, and catalase and cytochrome oxidase positive. Growth occurs from 2 °C to 30 °C. Optimal growth temperature occurs at 15 – 20 °C. The pH range for growth is 5.0 – 9.0 with an optimum at 7.0. The species requires NaCl to grow and grows at NaCl concentrations of 1 – 3 % with an optimum between 1 and 2 %. It utilizes D-xylose, D-galactose, D-glucose, D-fructose, N-acetylglucosamine, D-cellobiose and D-maltose, and it hydrolyses potassium 5-ketogluconate and esculin. Acid is produced from glucose by fermentation, and the species degrades starch and produces catalase, β-galactosidase, and cytochrome oxidase. It reduces nitrate but does not reduce nitrite. It is negative for production of indole, arginine dehydrolase, urease and gelatin hydrolase. The cellular fatty acid profile is dominated by C_16:0_, C*_16:1_^ω7c/^* C*_16:1_^ω7c/^* and C*_14:0_*^3OH^ fatty acids.

The type strain MME1^T^ (= DSM 118464^T^ =LMG 33722^T^) was isolated from estuarine sediment collected in a coastal area where the Nivå river meets the Baltic Sea, named Nivå Bugt Strandenge Bird Sanctuary, Denmark, in the spring of 2021. The genomic G+C content of the type strain was 39.5 mol%.

## FUNDING INFORMATION

The work was supported by the Novo Nordisk Foundation [NNF21OC0070749], the Independent Research Fund Denmark [grant DFF – 8048-00035B], and the National Danish Research Foundation [DNRF137].

## ACKNOWLEDGMENTS

We wish to thank MME group students Mathilde Engelhardt Petersen, Asger Bille, Caroline Toft Stripp, and Esther Fhær Iversen for preliminary work with the isolation and characterization of this species. We wish to thank Jette Melchiorsen and Haura Al-joubouri for technical assistance. We also thank the colleagues from the DSMZ and the Belgian Coordinated Collections of Microorganisms (BCCM) for their support in strain deposit.

## CONFLICT OF INTEREST

The author declares no conflicts of interest.

## Notes

### Competing Interest Statement

The authors have declared no competing interest.

