## Supplementary figure 1 for "*Psychromonas aestuarii* sp. nov., a novel bacterial species isolated from estuarine surface sediment"

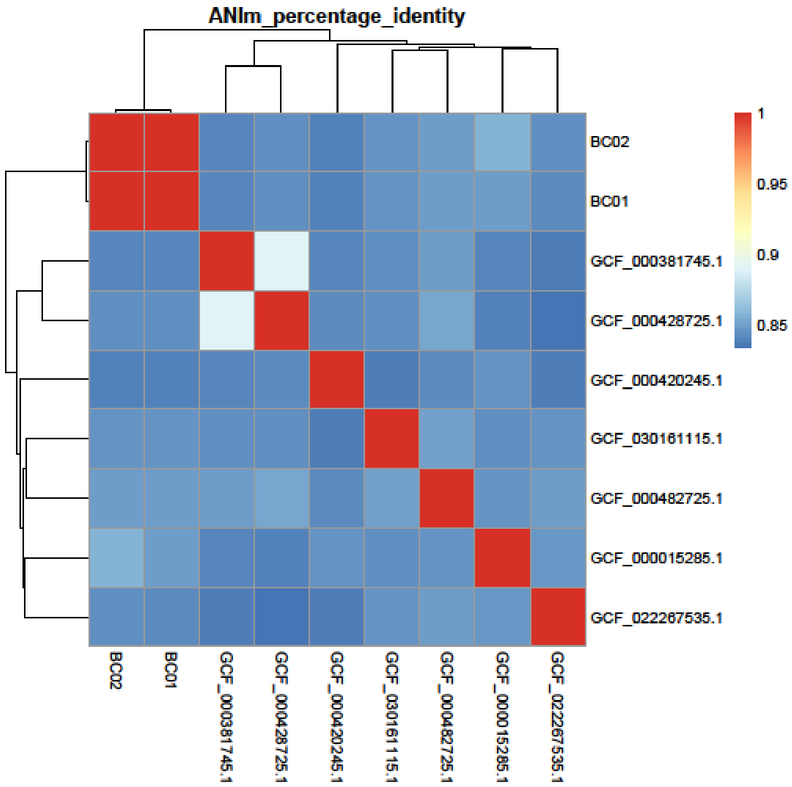


**Fig S1**. ANI percentage identity. *Psychromonas aestuarii* MME2 (BC02 or GCF_040566515.1); *Psychromonas aestuarii* MME1(BC01 or GCF_041080865.1); *Psychromonas ossibalaenae* ATCC BAA-1528 JMM 0738$^{T}$ (GCF_000381745.1); *Psychromonas aquimarina* ATCC BAA 1526$^{T}$ (GCF_000428725.1); *Psychromonas hadalis* ATCC BAA-638$^{T}$ (GCF_000420245.1); *Psychromonas marina* NBRC 103166$^{T}$ (GCF_030161115.1); *Psychromonas arctica* DSM 14288$^{T}$ (GCF_000482725.1); *Psychromonas ingrahamii* 37$^{T}$ (GCF_000015285.1); *Psychromonas antarctica* DSM 10704$^{T}$ (GCF_022267535.1).
